# Investigating the role of stomatal dynamics on agronomic traits using a *slac1-2 Zea mays* mutant

**DOI:** 10.1101/2025.01.21.634166

**Authors:** Robert J. Twohey, Clay G. Christenson, Catherine Li, Harel Bacher, Sebastian Calleja, Bryan Pastor, Marjorie Hanneman, Liam Wickes-Do, Michael A. Gore, Duke Pauli, Stephen P. Moose, Anthony J. Studer

**Affiliations:** Department of Crop Sciences, University of Illinois Urbana-Champaign, Urbana, IL 61801, USA; The School of Plant Sciences, University of Arizona, Tucson, AZ 85721, USA; Plant Breeding and Genetics Section, School of Integrative Plant Science, Cornell University, Ithaca, NY 14853, USA

**Keywords:** *Zea mays*, *slac1-2*, stomatal dynamics, field trials, leaf gas exchange, grain yield, canopy temperature, nitrogen utilization, and δ^13^C_leaf_

## Abstract

The production of staple food crops is becoming increasingly difficult due to the amount of freshwater needed to realize food security for a growing global population. Climate projections show hotter and drier growing seasons in traditionally productive agricultural regions, which will increase the demand for limited water resources. Thus, strategies to improve water use efficiency will only become more important for sustainable agriculture. At the leaf level, stomatal conductance has a large influence on transpirational water loss and therefore water use efficiency. The anion channel SLAC1 has been shown in several plant species to function as the primary mechanism for stomatal closure, which allows stomatal aperture to change dynamically in response to environmental stimuli. Given that *slac1* is a single gene that can significantly alter stomatal conductance, it has the potential to serve as a control point for improving water use efficiency. Here we fully characterize a *Zea mays slac1-2* mutant in multiple field environments. Interestingly, homozygous *slac1-2* hybrids did not show improved net CO_2_ assimilation or increased grain yield despite having greater stomatal conductance compared to a wild-type hybrid. Net CO_2_ assimilation and grain yield were either lower or similar in *slac1-2* compared to wild-type across environments. Furthermore, the *slac1-2* hybrid did not have increased nitrogen uptake. These results suggest that the C_4_ carbon concentrating mechanism removes any stomatal conductance limitations to CO_2_ assimilation, even in highly productive wild-type *Z. mays* hybrids. Because the *slac1-2* hybrids eliminate stomatal conductance as a major variable for modulating water loss, future studies will be able to investigate alternate regulators of plant water potential to identify novel mechanisms for increasing water use efficiency.

## Introduction

The production of staple food crops currently demands a large amount of water. It is estimated that current agricultural practices account for 70-75% of human freshwater consumption (McDaniel *et al*., 2017; Wallace, 2000). Climate projections predict a hotter and drier future, which would further increase agricultural water demand. The 2024 global average surface temperature was 1.6°C above pre-industrial levels and is expected to continue to rise through 2040 (Copernicus Climate Change Service, 2024; IPPC, 2023). Precipitation patterns have also become more sporadic, increasing the occurrence and severity of droughts (Gamelin *et al*., 2022). Together, these climate variables drive the observed increases in atmospheric vapor pressure deficit (VPD), defined as the difference between the amount of moisture in the air and its saturation point. The drastic rise in VPD during growing seasons is significantly impacting crop productivity, and rectifying its effects remains challenging (López *et al*., 2021; Novick *et al*., 2024). Combined precipitation deficits and high VPD can result in flash droughts during the growing seasons, which are harder to predict and control through traditional management practices (Christian *et al*., 2021). While current irrigation practices can suppress crop loss due to drought, studies show that increasing irrigation to combat climate change will not be a sustainable management strategy (McDonald and Girvetz, 2013). Therefore, efforts targeting biological processes to better understand and improve water use efficiency (*WUE*) strategies in our major crops are needed (Leakey *et al*., 2019).

Leaf level *WUE* is greatly influenced by stomatal regulation of gas exchange between the atmosphere and the intercellular airspace of the leaf. Through changes in aperture, the ratio between CO_2_ assimilation (*A*_net_) necessary for photosynthesis and water loss via transpiration (*E*) can dynamically change in response to environmental stimuli (Wong et al., 1979; Assmann and Jegla, 2016; Lawson and Vialet-Chabrand, 2019). The precise control of CO_2_ and water flux is vital for maintaining productive and efficient plants (Farquhar and Sharkey, 1982; Leakey *et al*., 2019). Vascular plants control stomatal conductance (*g*_s_) by altering stomatal aperture using changes in the turgor pressure of surrounding guard cells (Keller *et al*., 1989). Stomatal response to environmental factors is highly dependent on biochemical signaling cascades and the speed of solute flux, as the movement of anions across the guard cell plasma membrane results in stomatal aperture changes (Mansfield, 1983; Lawson and Blatt, 2014).

One of the main guard cell-specific anion channels, SLAC1 (Slow Anion Channel-Associated 1), was originally characterized in *Arabidopsis thaliana* (Negi *et al*., 2008). A loss of function mutant in *At*SLAC1 resulted in over-accumulation of osmoregulatory anions and hyposensitive stomatal response (Negi *et al*., 2008; Vahisalu *et al*., 2008). Upstream signaling pathways activate *slac1* through the phosphorylation at its N terminus, allowing for the efflux of anions out of the guard cells and stomatal closure (Geiger *et al*., 2009; Lee *et al*., 2009). Multiple environmental stimuli alter slow anion channels such as light, humidity, atmospheric CO_2_, reactive oxygen species, and abscisic acid (ABA) drought response (Pei *et al*., 1997; Pei *et al*., 2000; Roelfsema and Hedrich, 2005; Negi *et al*., 2008; Vahisalu *et al*., 2008). Rather than all stimuli independently altering SLAC1 through isolated signaling pathways, these environmental cues converge to fine-tune a core ABA signaling pathway that ultimately phosphorylates SLAC1 (Hedrich and Geiger, 2017).

In addition to *A. thaliana*, SLAC1 orthologs have been studied in crop species such as *Hordeum vulgare* (Liu *et al*., 2014), *Gossypium herbaceum* (Ren *et al*., 2021), *Oryza sativa* (Kusumi *et al*., 2017), *Solanum tuberosum* (He *et al*., 2025), and *Zea mays* (Qi *et al*., 2018). In this study, we use a previously identified *Z. mays slac1-2* UniformMu transposable element insertion (mu1037824), which creates a null mutation. *Z. mays* contains a single gene copy of *slac1* with 67.4% protein homology to the *A. thaliana* ortholog (Goodstein *et al*., 2012). Initial characterization by Qi *et al*. (2018) found *slac1-2* to have similar hyposensitive stomatal responses to changing environmental conditions, and to be highly nitrate-selective with less permeability to other anions compared to *At*SLAC1. Other work in *Z. mays* showed multiple protein kinases (ZmCPK35 and ZmCPK37) and upstream phosphatases (ZmPP84) to activate *slac1* through signaling cascades. However, a complete understanding of stomatal signaling pathways in *Z. mays* is still lacking and its similarity to *A. thaliana* is not fully known (Li *et al*., 2022; Guo *et al*., 2023).

In C_3_ species, the tight connection between *g*_s_ and *A*_net_ have long been known. Many studies have shown increased atmospheric CO_2_ levels or increased *g*_s_ leading to increases in *A*_net_ and yield (von Caemmerer and Evans, 2010; Lawson *et al*., 2011; Ainsworth and Long, 2012; Kusumi *et al*., 2012). Multiple *slac1* mutants in C_3_ species have been characterized to better understand the dynamic relationship between *slac1* activation, *g*_s_, *A*_net_, and yield. In barley, positive correlations were found between HvSLAC1 expression and salt tolerance, indicating that increased stomatal closure improved salt tolerance and yield (Liu *et al*., 2014). Alternatively, a greenhouse grown rice *slac1* mutant with higher *g*_s_ resulted in higher photosynthetic rates (Kusumi *et al*., 2012). The relationship between *g*_s_ and *A*_net_ is thought to be weaker in C_4_ species due to the presence of a CO_2_ concentrating mechanism, which saturates CO_2_ around Rubisco in bundle sheath cells. If Rubisco is always CO_2_ saturated, it eliminates the opportunity to realize increased photosynthesis and yield by increasing *g*_s_. In this study, *slac1-2* hybrids were used to further determine if *Z. mays* productivity is limited by the stomatal regulation of *g*_s_.

Here, we fully characterize an inbred *slac1-2* mutant in a dynamic field environment. *Z. mays* hybrids homozygous for *slac1-2* were also developed and evaluated at multiple field locations representing contrasting environments. These trials allowed us to directly test the performance of hybrids with increased *g*_s_ and lack of dynamic stomatal response across variable field environments. Collecting leaf gas exchange, grain yield, and nitrogen utilization data from these field trials furthers our understanding of the relationships between physiological traits and agronomic performance in a major C_4_ field crop and indicates strategies for improving *WUE*.

## Material & Methods

### Germplasm

The *slac1-2* UniformMu insertion line (mu1037824 in stock UFMu 04043) was acquired from the USDA Maize Genetic Stock Center located at the University of Illinois (Portwood *et al*., 2019). Segregating seed was selfed in the greenhouse and field for multiple generations in 2017 to produce homozygous stocks of *slac1-2* and its corresponding wild-type in a W22 background. During each generation, genetic screens were used to produce non-segregating Mu stocks. Tissue was collected from one week old seedlings and immediately placed in liquid nitrogen. DNA extractions were done following a CTAB protocol as described in (Kolbe *et al*., 2018). A forward AJS537 (AGCAGAGAGAAGACTTGCGG) and reverse AJS538 (TGGACGGGGAAACTTTGTAG) primer were designed to flank the transposable element insertion (mu1037824). A previously designed Mu specific primer AJS516 (GCCTCYATTTCGTCGAATCCS) was used that aligns with the inverted terminal repeats located on each end of the transposable element (Settles *et al*., 2004). Mu1037824 insertion was confirmed using genic-genic (AJS537/538) and genic-mutator (AJS537/516) primer sets with a Phire Hot Start II (#F-122L) PCR reaction following manufacturers recommendations.

To produce the B73 X H99 *slac1-2* two-way hybrid, *slac1-2* was first crossed with the H99 and B73 inbred lines. Each line was then backcrossed multiple generations and selfed to produce BC_5_S_1_ seed. During backcrossing, the AJS537/516 primer set was used to confirm that the *slac1-2* allele was retained. Homozygous B73 X H99 *slac1-2* and its corresponding B73 X H99 wild-type hybrids were then planted at each field site.

### Plant Growth

Greenhouse grown *slac1-2* and wild-type inbreds were planted in 50 well plug trays (T.O. Plastics #720568C) with a 1:1 mixture of Sun Gro Sunshine LC1 and a general purpose 1:1:1 – soil : peat : perlite mix with 2.3 kg of dolomitic lime, 0.9 kg of 0-46-0, 1.4 kg of gypsum, and 0.9 kg of Epsom per cubic yard. Plants remained in flats for 2 weeks and then were transplanted into a greenhouse ground bed. Plants were fertilized with 15-5-15 CalMag at a concentration of 300 ppm weekly and Sprint 330 chelated iron at 30 ppm was applied once to all plants around v5.

During the 2021 field season, homozygous *slac1-2* and wild-type inbreds were planted at the Crop Sciences Research and Education Center located in Urbana, Illinois under irrigation. Twenty kernels were planted in a 3.7 m row with 0.8 m spacing between rows and 0.9 m alleys. Nitrogen was applied to the field at a rate of 156.9 kg/ha before planting. Two rows of each genotype were planted, and six representative plants were chosen for analysis excluding end plants.

The B73 X H99 *slac1-2* hybrids were grown in 2022, 2023, and 2024. There were four different field locations. Illinois location 1 was the same irrigated field described for inbred measurements and Illinois location 2 was a non-irrigated field less than 1 km away. Both Illinois hybrid trials were planted in 2022 and 2023 containing thirty-five kernels planted in a 5.2 m row with 0.8 m spacing between rows and 0.9 m alleys. Illinois location 1 had nitrogen applied at a rate of 156.9 kg/ha before planting while Illinois location 2 had variable nitrogen rates that are described in the R6 sampling methods. The New York field location was located at Cornell University’s Musgrave Research Farm in Aurora, NY, trials were run in 2022 and 2023. The New York trial contained thirty-five kernels planted in 5.2 m two-row plots with 0.8 m spacing between rows and 0.9 m alleys. The New York field had 177.7 L/ha of liquid 12.5-25-0 band applied pre-planting and 318 L/ha of 30%UAN side-dressed at v5-v6. The Arizona trial was located at the University of Arizona’s Maricopa Agricultural Center in Maricopa, Arizona. The Arizona trial contained forty kernels planted in a 5.4 m row with 1 m spacing between rows and 0.7 m alleys. Total nitrogen application in Maricopa was approximately 261.2 kg/ha across the growing season. Approximately 37 kg/ha were applied preplant (DOY 71) via granular fertilizer with approximately 224.2 kg/ha applied in-season with four splits of 56 kg/ha occurring on DOY 117, 127, 139, and 155 delivered as UAN 32 applied through flood irrigation.

### Thermal Imaging

The infrared imaging was taken in 2021 at the Illinois location 1 field site from a ladder approximately 3 m above ground level using a FLIR ThermaCAM T-400 Wes Camera (SN: 345001389) when plants were at v10. The FLIR thermal studio standard software (Version: 2.0.26) was used to capture n = 4 random but representative sampling locations on the canopy surface for each genotype. The average leaf temperature was calculated within the standardized area and used as a biological replicate.

### δ^13^C_leaf_ Isotope Collection and Analysis

All δ^13^C_leaf_ sample collection was performed as described in Twohey III *et al*. (2019). During the 2018 greenhouse and 2021 Illinois location 1 field season, δ^13^C_leaf_ samples were collected from healthy, uppermost fully expanded, v10-12 leaves. Leaf samples were collected from individual plants for a total of n = 4 greenhouse samples and n = 6 field samples. Each sample contained 6-12 tissue punches from each side of the midrib totaling 12-24 leaf punches per sample. Leaf tissue samples were then dried at 65°C for 7 d, ground, and stored in a desiccation cabinet until isotope analysis.

Leaf stable carbon isotope analysis was run as described in Twohey III *et al*. (2019). The samples were run at the University of Illinois through a Costech Instruments elemental combustion system, and then a Delta V Advantage mass spectrometer to determine δ^13^C_leaf_ values. The instrument precision has been shown to be ±0.2‰ when measuring δ^13^C. Vienna Peedee Belemnite was used as a calibration standard.

### Stomatal Density

Three tissue samples (5 x 3cm) were collected from each greenhouse grown plant. Samples were immediately placed in liquid nitrogen then transferred to a -80°C freezer. Leaf samples were thawed and placed on microscope slides using double sided tape before imaging. A μsurf explorer optical topometer with a 20×/0.60 objective lens (image size 0.8 × 0.8 mm^2^) was used for imaging (Nanofocus, Oberhausen, Germany). Total number of stomata were counted in each image. Three images were taken on each side of the leaf sample and averaged to produce one biological replicate (n=12).

### Li-6800 Steady-State and Response Curve Measurements

The inbred *slac1-2* measurements were taken using a LI-6800 photosynthesis system (LI-COR Biosciences, Lincoln, NE, USA) with a 3 x 3cm chamber and LED light source. Chamber temperature was set to 29°C during the 30 min. acclimation stage, and then leaf temperature was held constant during measurements. Humidity was controlled to maintain a leaf VPD of 1.5 kPa. Light was set constant at 2,000 μmol m⁻² s⁻¹. Individual plants were measured as biological replicates (n=6).

Gas exchange measurements started 1 h after daybreak and ended by 14:00 the same day. For each plant measured, the leaf was clamped and remained under constant conditions at 400 ppm CO_2_ for at least 30 min. When steady state was achieved, data collection began, and a measurement was taken every 10 s throughout the time course. Data points were logged at 400 ppm CO_2_ for 30 min. Then the CO_2_ concentration of the chamber was increased to 800 ppm and data points were logged for 30 min. allowing for the observation of stomatal closure rates and steady state values under a high CO_2_ concentration. The final CO_2_ change was a decrease to 100 ppm to observe stomatal opening rates. Data points were logged for 1 h at 100 ppm CO_2_ due to a longer acclimation time necessary to reach steady state.

### Li-6800 survey measurements

Survey measurements were taken on sunny days between 11:00 and 13:00. The hybrid *slac1-2* measurements were taken using a Li-6800 gas exchange system with a 3 cm^2^ chamber and LED light source. Flow rate was set to 500 µmol s^-1^. The remaining environmental conditions were set to replicate atmospheric CO_2_, temperature, light, and humidity at the time of measurement. Plants were measured on the uppermost fully expanded leaf once v12 was reached. Once a leaf was clamped, a log was recorded after all environmental conditions reached steady state. Biological replicates varied based on location (Illinois location 1 n=12, Arizona n=24).

### Grain Yield Data

To measure grain yield data, total ears from five plants were collected from each field trial row making sure to exclude end plants. Biological replicates varied based on location (Illinois location 1 n=12, Illinois location 2 n=6, New York n=8, Arizona n=12). The ears were dried for at least seven days at approximately 65°C. Each row was then shelled, and grain weight and grain moisture were collected. The grain weight and moisture were used to calculate grain yield at a normalized moisture of 15.5%. When grain yield data was used to calculate total Mg/ha, differences in row spacing and density were accounted for.

### R6 Sampling and Analysis

Nitrogen uptake and utilization were determined using whole shoots sampled at physiological maturity (R6) when at least 50% of the plants showed a visible black layer at the base of the kernel. Biomass and nitrogen samples were collected and measured as described in (Cheng *et al*., 2021). Biomass nitrogen combustion analysis with a Fisons EA-1108 N elemental analyzer and grain nitrogen analysis using a near-infared reflectance spectroscopy on a Perten DA7200 analyzer was completed at the University of Illinois. Biological replicates varied for 2022 and 2023 (Table 2).

### Statistical analysis

Significant differences between *slac1-2* and wild-type plants were tested using the (t.test) function (version 3.6.2) in R (R Core Team (2021)) by running a Welch’s *t*-test.

## Results

### Canopy thermal imaging

Although the effect of mutations in *slac1* have been documented at the leaf level, less is known about the extent to which the mutation alters canopy traits in the field. Thermal images were taken to determine if *slac1-2* leaf temperature differences could be observed at a field canopy level. The *slac1-2* inbred showed an average canopy temperature of 27.58 ± 0.19 °C. The wild-type comparator had a significantly higher canopy temperature (28.83 ± 0.08 °C) compared to *slac1-2 (P <* 0.001, *t*-test, Fig. 1). These results indicate that leaf level temperature scales to the canopy level in field grown *slac1-2* mutants.

**Figure 1.**
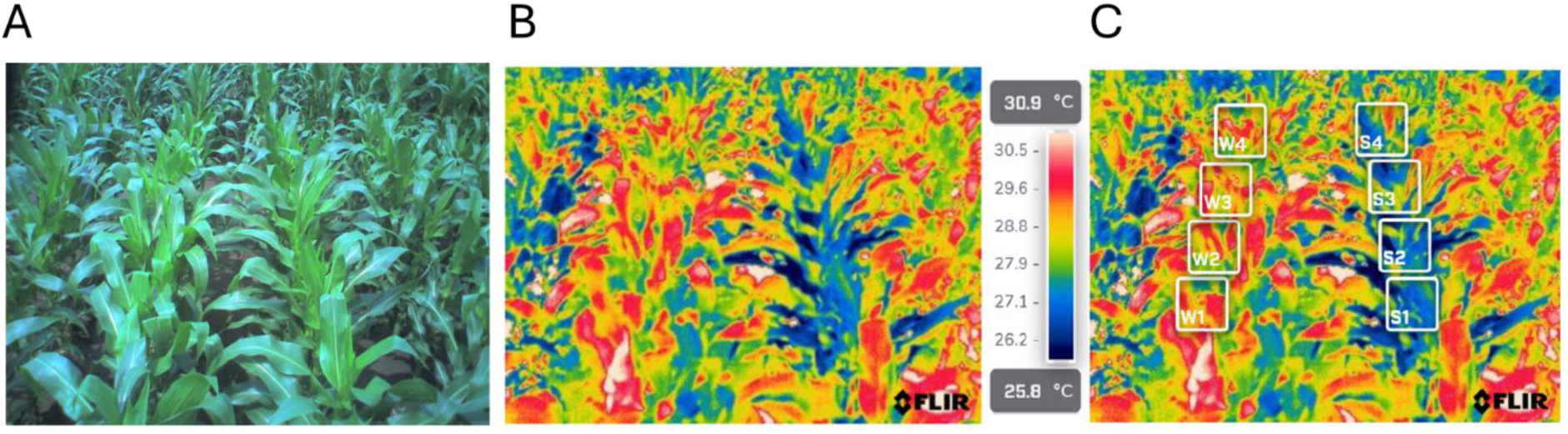
Infrared image taken with a ThermaCAM T400 Camera. **A**) RGB photo taken in 2021 field trial of *slac1-2* and wild-type rows, **B**) Thermal image of A, **C**) Temperature analysis. The boxes in panel C represent technical reps of canopy temperature. Average temperature was calculated for the area within each square, wild-type (28.83 ± 0.08 °C) (Left, W1-4), *slac1-2* (27.58 ± 0.19 °C) (Right, S1-4).

### Leaf isotope composition analysis of field and greenhouse grown slac1-2 inbreds

The *slac1-2* inbred and its comparative wild-type were planted during the 2018 greenhouse and 2021 field season to determine how the *slac1-2* stomatal phenotype affects leaf isotope composition. In the greenhouse, *slac1-2* showed significantly lower percent carbon (%C) values compared to wild-type, (*P* < 0.01, *t*-test, Table 1) while percent nitrogen (%N) was not significantly different (Table 1). Significantly less negative δ^13^C_leaf_ values were observed in greenhouse grown *slac1-2* plants compared to wild-type (*P* < 0.01, *t*-test, Table 1). In the field, *slac1-2* had significantly lower %C and significantly higher %N compared to wild-type (*P* < 0.05 and *P* < 0.01 respectively, *t*-test, Table 1). Similar to the greenhouse, field grown *slac1-2* also showed less negative δ^13^C_leaf_ values compared to wild-type (*P* < 0.001, *t*-test, Table 1).

**Table 1.** Stable carbon isotope values of greenhouse and field grown *slac1-2* and wild-type W22 inbred *Z. mays*.

|  | Greenhouse Grown |  | Field Grown |  |
| --- | --- | --- | --- | --- |
|  | Wild-type | *slac1-2* | Wild-type | *slac1-2* |
| %C | 45.75±0.12** | 44.82±0.17** | 47.76±0.07* | 47.55±0.03* |
| δ<sup>13</sup>C | -13.52±0.07** | -12.48±0.18** | -12.98±0.06*** | -12.47±0.05*** |
| %N | 4.47±0.08 | 3.88±0.28 | 4.12±0.06** | 4.4±0.05** |

\*= $P < 0.05$ , \*\*= $P < 0.01$ , and \*\*\* $P < 0.001$ indicates significant threshold cutoffs for *t*-test comparisons of *slac1-2* and wild-type for each trait blocked by grow out.
Percent carbon = %C, stable carbon isotope ratio = δ<sup>13</sup>C, percent nitrogen = %N, stable nitrogen isotope ratio = δ<sup>15</sup>N

### Stomatal density

Stomatal density was measured in *slac1-2* inbred and wild-type plants, grown in the 2018 greenhouse, to determine if gas exchange differences observed in *slac1-2* were solely due to dynamic aperture differences. No significant differences in stomatal density were observed between *slac1-2* and wild-type plants. Average abaxial density was 52.36±1.67 in *slac1-2* and 54.58±1.43 in wild-type (*P* = 0.32, *t*-test, Sup. Fig. 1). Average adaxial density was 42.69±1.06 in *slac1-2* and 43.14±1.35 in wild-type (*P* = 0.80, *t*-test, Sup. Fig. 1). Therefore, the observed gas exchange differences were due to stomatal aperture and not stomatal density differences between *slac1-2* mutant and wild-type plants.

### Field measured slac1-2 inbred response to CO_2_

Previously, leaf level gas exchange was measured to determine *slac1-2* response to CO_2_ in a controlled environment with rapid (<20 minute) response curves (Qi *et al*., 2018). To capture the full dynamic response, extended time-course measurements were performed that allowed plants to reach a steady-state at each CO_2_ concentration. Furthermore, the *slac1-2* response to changes in CO_2_ concentration was measured using field grown plants. Leaf *g*_s_ was measured at three different atmospheric CO_2_ concentrations. During the 400ppm and 800ppm CO_2_ steady state, *slac1-2* had significantly higher *g*_s_ compared to wild-type. (*P* < 0.01, *t*-test, Fig. 2A, Sup. Table 1). At a 100ppm steady state, no significant differences were observed. The same trend was seen for transpiration (*E*) where *slac1-2* rates were significantly higher compared to wild-type at 400ppm and 800ppm steady states (*P* < 0.01, *t*-test, Sup. Table 1). No significant difference in *A*_net_ was observed between *slac1-2* and wild-type across all CO_2_ concentration steady states (Fig. 2B, Sup. Table 1). During the CO_2_ stepwise response measurements, *slac1-2* had minimal response to increased or decreased CO_2_ concentrations compared to wild-type (Fig. 2C).

**Figure 2.**
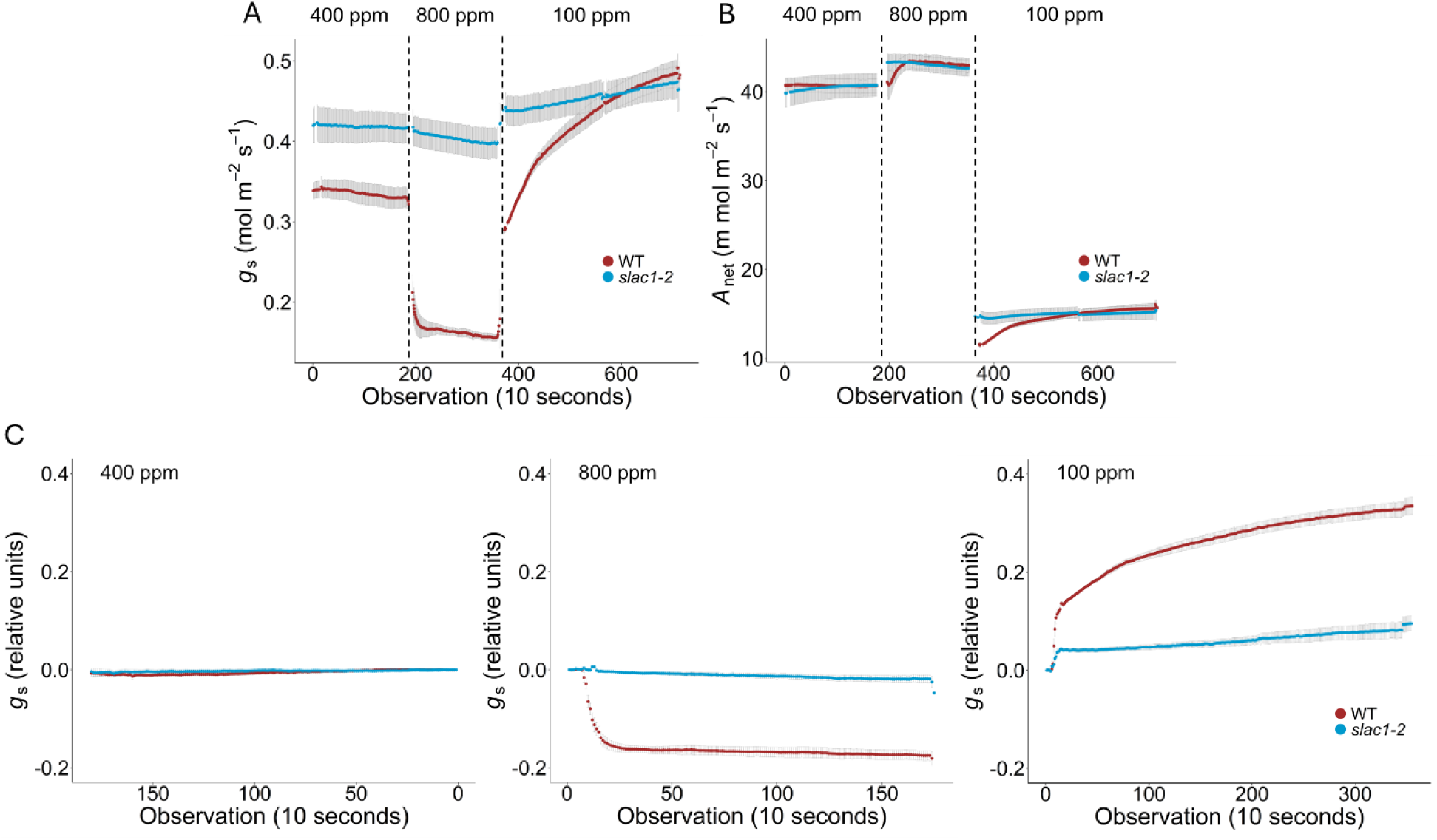
During the 2021 Illinois location 1 field season, stomatal response to atmospheric CO_2_ was measured using gas exchange in a *slac1-2* and wild-type W22 background. A) Average *g*_s_ level ± standard error at ten second intervals for *slac1-2* (blue) and wild-type (WT, red) during three different CO_2_ concentrations defined at the top of each graph (400ppm, 800ppm, and 100ppm). Vertical dashed lines indicate change in CO_2_ concentration. B) Average *A*_net_ ± standard error at ten second intervals for *slac1-2* and wild-type during three different CO_2_ concentrations. C) Average *g*_s_ ± standard error is normalized to the final point collected during the previous CO_2_ concentration. Initial steady state values are normalized to the final 400 ppm data point.

### Hybrid slac1-2 mid-day gas exchange measurements

Mid-day gas exchange measurements were taken using the Li-6800 survey method. At the 2022 Illinois location 1 field site, significantly higher mid-day rates of *g*_s_ were observed in the *slac1-2* hybrid compared to wild-type (*P* < 0.05, *t*-test, Fig. 3A). Due to higher *g*_s_ rates, significantly higher *E* and intercellular CO_2_ (*C*_i_) values were also observed (*P* < 0.05 and *P* < 0.01 respectively, *t*-test, Fig. 2B and D). Higher *g*_s_ rates resulted in significantly lower leaf temperature in *slac1-2* compared to wild-type (*P* < 0.01, *t*-test, Fig. 3F). Although higher *C*_i_ levels were observed, there was no significant difference in *A*_net_ between the *slac1-2* and wild-type hybrid (Fig. 2E). Due to the observed increase in *g*_s_ with no benefit to *A*_net_, intrinsic *WUE* (*iWUE*, A_net_/*g*_s_) was significantly lower in the *slac1-2* hybrid compared to wild-type (*P* < 0.01, *t*-test, Fig. 2C). Mid-day gas exchange measurements taken during the 2023 Illinois location 1 and 2024 Arizona field seasons showed no significant differences in *g*_s_, *E*, *C*_i_, and *A*_net_/*g*_s_ when the *slac1-2* hybrid was compared to its respective wild-type hybrid (Fig. 3). Significantly lower *A*_net_ values were observed in the *slac1-2* hybrid during the 2023 Illinois location 1 mid-day measurements (*P* < 0.05, *t*-test, Fig. 2E).

**Figure 3.**
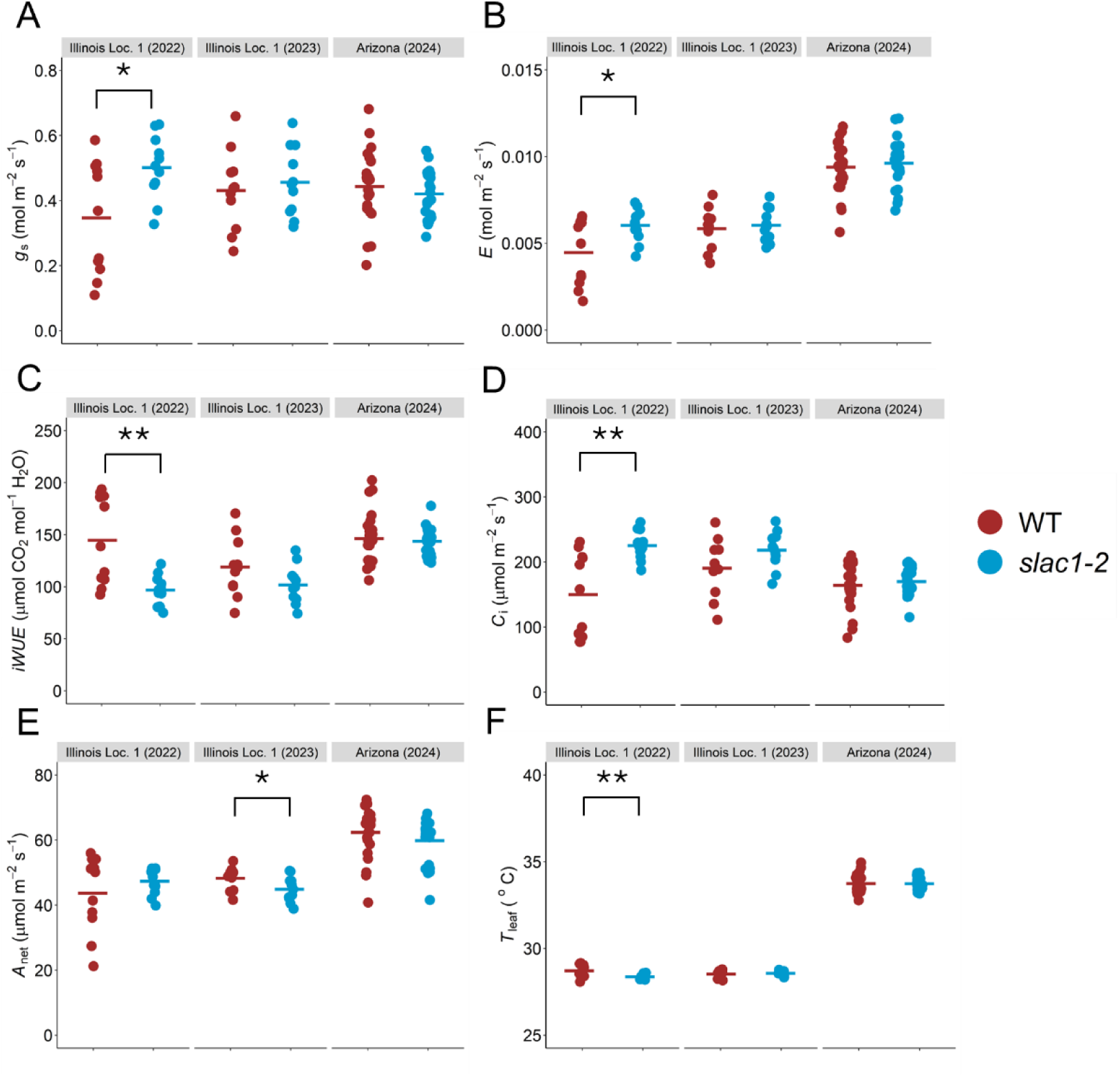
Survey measurements were taken with a Li-6800 during the 2022 and 2023 Illinois location 1 and 2024 Arizona field seasons on a sunny day between 11:00 and 12:00. A) Stomatal Conductance (*g*_s_), B) Transpiration (*E*), C) Intrinsic water use efficiency (*iWUE, A*_net_/*g*_s_), D) Intercellular CO_2_ (*C*_i_), E) Net CO_2_ Assimilation (*A*_net_), F) Leaf Temperature (*T*_leaf_). Points represent individual replicates, and the solid dash represents the average for each trait. Significant differences between *slac1-2* (blue) and wild-type (WT, red) are denoted by * = *P* < 0.05 or ** = *P* < 0.01 (*t-*test).

To investigate the relationship between *g*_s_ and *A*_net_ in *Z. mays*, *A/C*_i_ curves were measured on *slac1-2* and wild-type hybrids during the 2022 field season. No differences in the rate of maximum carboxylation capacity of phosphoenolpyruvate (*V*_pmax_) was observed between the two genotypes. For *slac1-2* and wild-type, average *V*_pmax_ was 68.02±2.82 and 80.20±4.66 μmol CO_2_ m^-2^ s^-1^ respectively (*P* > 0.05, *t*-test, Fig. 4). No difference in CO_2_ saturated photosynthetic capacity (*V*_max_) was observed between *slac1-2* and wild-type, with an average *V*_max_ of 51.97±1.02 and 53.71±1.37 μmol CO_2_ m^-2^ s^-1^ respectively (*P* > 0.05, *t*-test, Fig. 4). The ratio between intercellular CO_2_ and atmospheric CO_2_ (*C*_i_/*C*_a_) was calculated at two points of the *A/C*_i_ curve when *C*_a_ was equal to 400ppm and 800ppm. At a *C*_a_ of 400ppm, *C*_i_/*C*_a_ was on average 0.49±0.02 and 0.42±0.03 for *slac1-2* and wild-type hybrids (*P* > 0.05, *t*-test, Fig. 4), respectively. When *C*_a_ reached 800ppm, significant differences were observed between *slac1-2* and wild-type plants, with averages of 0.70±0.01 and 0.39±0.03 respectively (*P* < 0.0001, *t*-test, Fig. 4).

**Figure 4.**
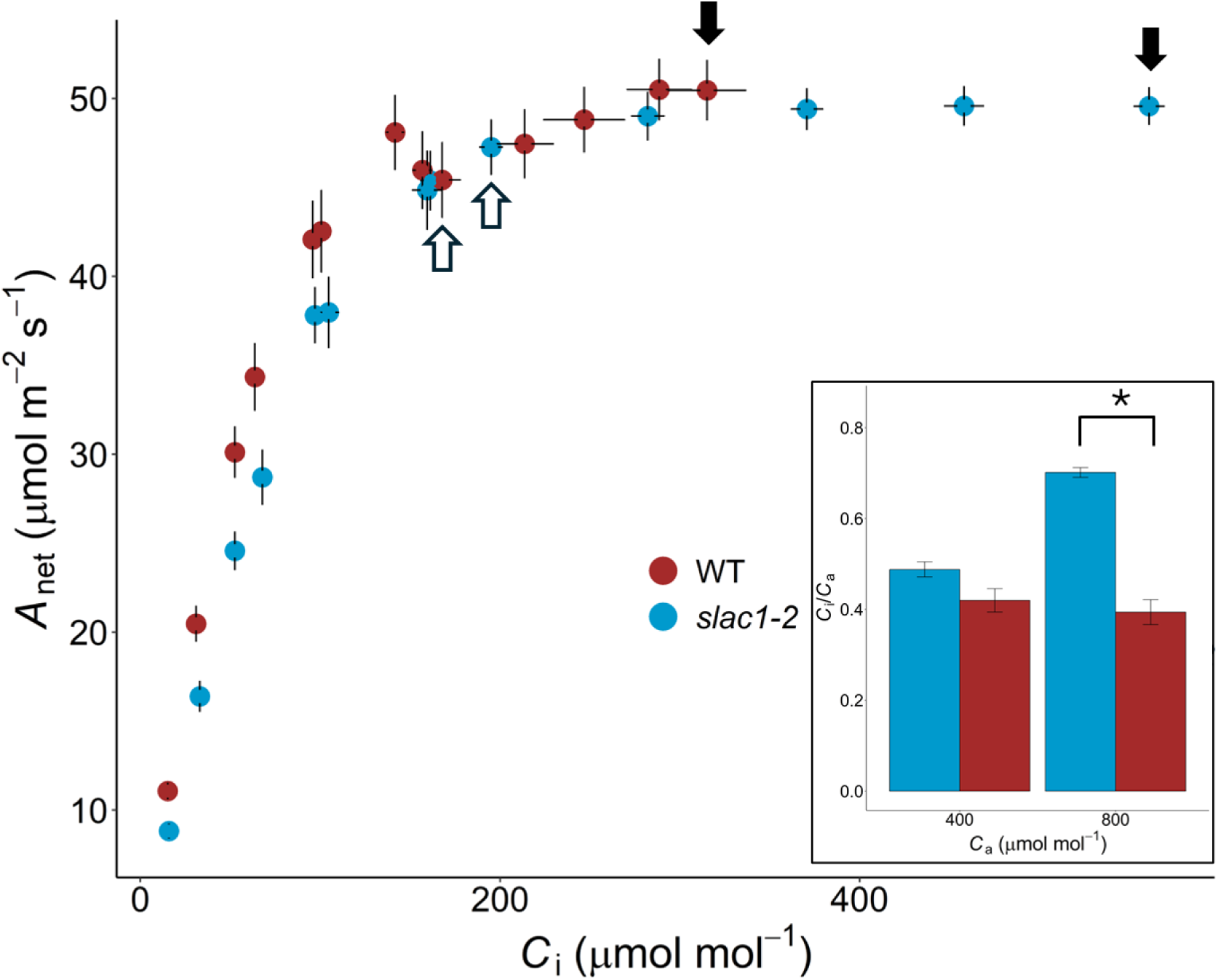
*A/Ci* curves were measured on *slac1-2* (blue) and wild-type (WT, red) hybrids during the 2022 Illinois location 1 field season. Net CO_2_ Assimilation (*A*_net_), Intercellular CO_2_ (*C*_i_), Ambient CO_2_ (*C*_a_). Black error bars represent standard error. Open and black filled arrows indicate the data used to calculate *C*_i_/*C*_a_ at a *C*_a_ of 400ppm and 800ppm CO_2_ respectively. The inset bar graph represents average *C*_a_/*C*_i_ ratios with error bars representing standard error (* = *P* < 0.05, *t*-test).

During the 2024 Arizona field season, a weather event increased cloud cover and caused a decrease in light intensity and a corresponding parabolic shaped VPD trend within the time span of four hours (7:00 – 11:00, Fig. 5). A Li-600 gas exchange system was used to measure *slac1-2* and wild-type hybrids throughout the morning to capture dynamic responses to changing VPD and light levels. During the first two hours of measurements (7:00 and 8:00) atmospheric VPD was 2.08 kPa and 1.98 kPa respectively. When the 9:00 measurements were taken, VPD had decreased to 1.34 kPa. Atmospheric VPD returned to 2.31 kPa before the final measurements were taken at 11:00. During this time span, the only significant difference between *slac1-2* and wild-type *g*_s_ was during the 9:00 measurement at a VPD of 1.34 kPa (*P* < 0.05., *t*-test, Fig. 5). These results demonstrate that the *slac1-2* hybrid lacked the ability to dynamically respond to environmental change (Fig. 5).

**Figure 5.**
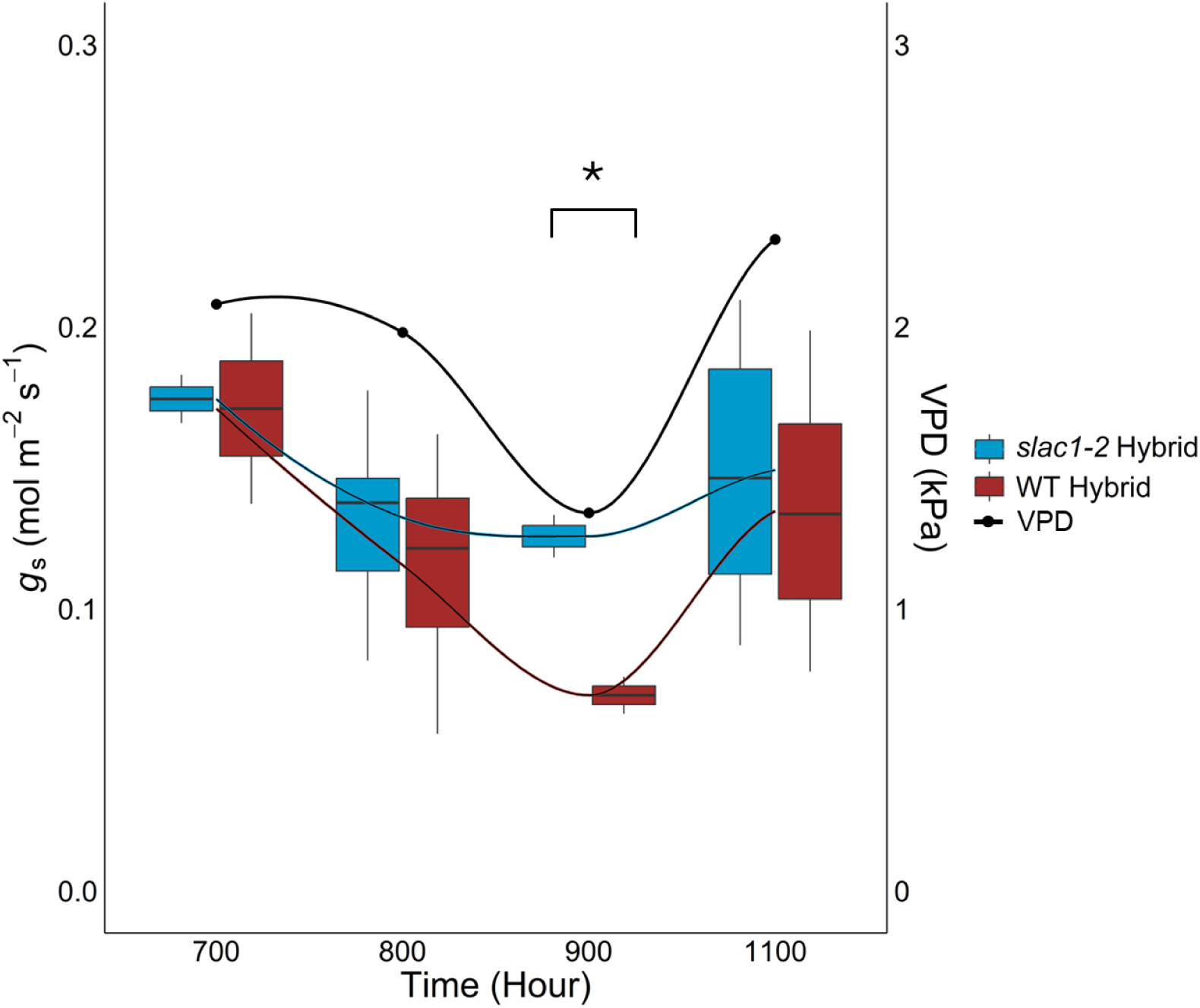
Gas exchange data was measured in the Arizona field trial using a Li-600 every hour from 7:00 to 11:00. The only significant difference between *slac1-2* (blue) and wild-type (WT, red) hybrids was during the 9:00 measurement (* = *P* < 0.05, *t*-test). The left y-axis and colored box plots represent the stomatal conductance (*g*_s_) values. The box plots show the first and third quartiles, with middle horizontal lines indicating the median and the whiskers showing the outliers. Smooth trend lines were added for each genotype. The right y-axis and black points with a smooth trend line represent vapor pressure deficit (VPD) values collected from the Maricopa field site weather station.

### Hybrid slac1-2 yield trials and nitrogen sampling

Grain yield was measured in seven different environments, across three years and four locations. Grain yield was found to be significantly different in four of the environments: Illinois location 1 2023, Illinois location 2 2022 and 2023, and Arizona 2024 (*P* < 0.01 for all, *t*-test, Fig. 6). While significant differences were observed between *slac1-2* and wild-type hybrids, *slac1-2* never out yielded wild-type. The *slac1-2* hybrid either had equal or lower grain yield compared to wild-type at all locations across all years (Fig. 6).

**Figure 6.**
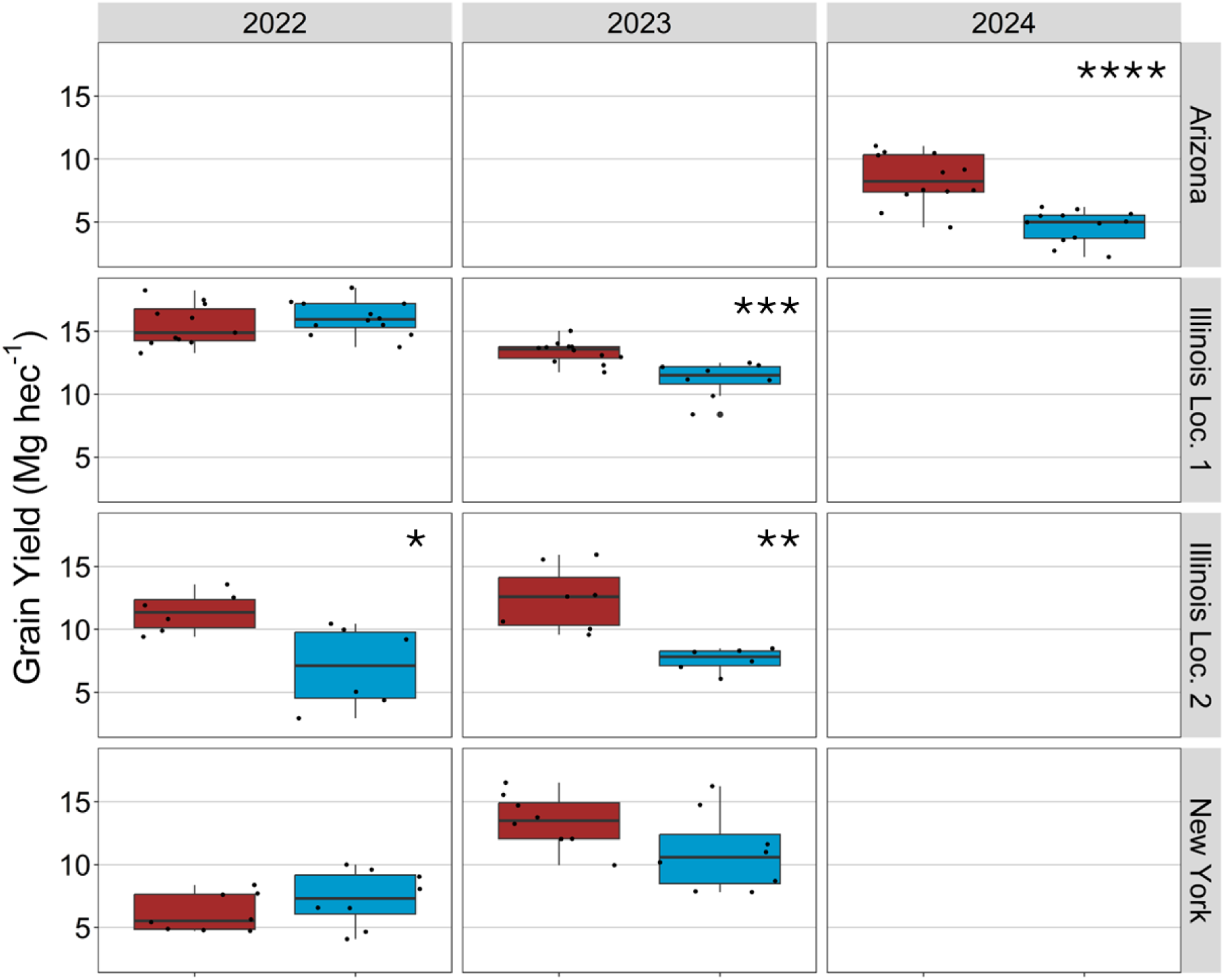
*slac1-2* (blue) and wild-type (red) hybrids were grown at multiple field locations to determine if stomatal control of CO_2_ uptake limits grain yield in *Z*. *mays. slac1-2* and wild-type grain yield data was normalized to 15.5% moisture and converted to megagrams per hectare (Mg hec^-1^) for all field sites. Field site years are denoted on the top x-axis and field locations are on the right y-axis. Significant differences between the two genotypes are denoted by * = *P* < 0.05, ** = *P* < 0.01, *** = *P* < 0.001, **** = *P* < 0.0001 (*t*-test).

In 2022 and 2023, R6 biomass and nitrogen content was measured under two nitrogen treatments (0 kg/ha and 225 kg/ha) at the Illinois location 2 field. Overall, trends showed that dry stalk biomass, cob biomass, dry stover biomass, and total biomass was lower in *slac1-2* compared to wild-type hybrids (*P* > 0.05, *t*-test, Table 2). In 2023, *slac1-2* plant height was significantly shorter than wild-type under 0 kg/ha nitrogen (*P* < 0.001, *t*-test) and 225 kg/ha nitrogen (*P* < 0.01, *t*-test, Supp. Table 2). Stalk nitrogen and stalk percent nitrogen was similar between the two genotypes, except stalk percent nitrogen was significantly greater in *slac1-2* compared to wild-type in the 2022 0 kg/ha treatment (*P* < 0.05, *t*-test, Table 2). Grain nitrogen was similar under 0 kg/ha applied nitrogen in both years and significantly lower in *slac1-2* compared to wild-type at 225 kg/ha nitrogen in 2022 and 2023 (*P* < 0.05 and *P* < 0.01 respectively, *t*-test, Table 2). Grain protein was significantly higher in *slac1-2* compared to wild-type across all years and nitrogen treatments (*P* < 0.05, *t*-test, Table 2).

**Table 2.**
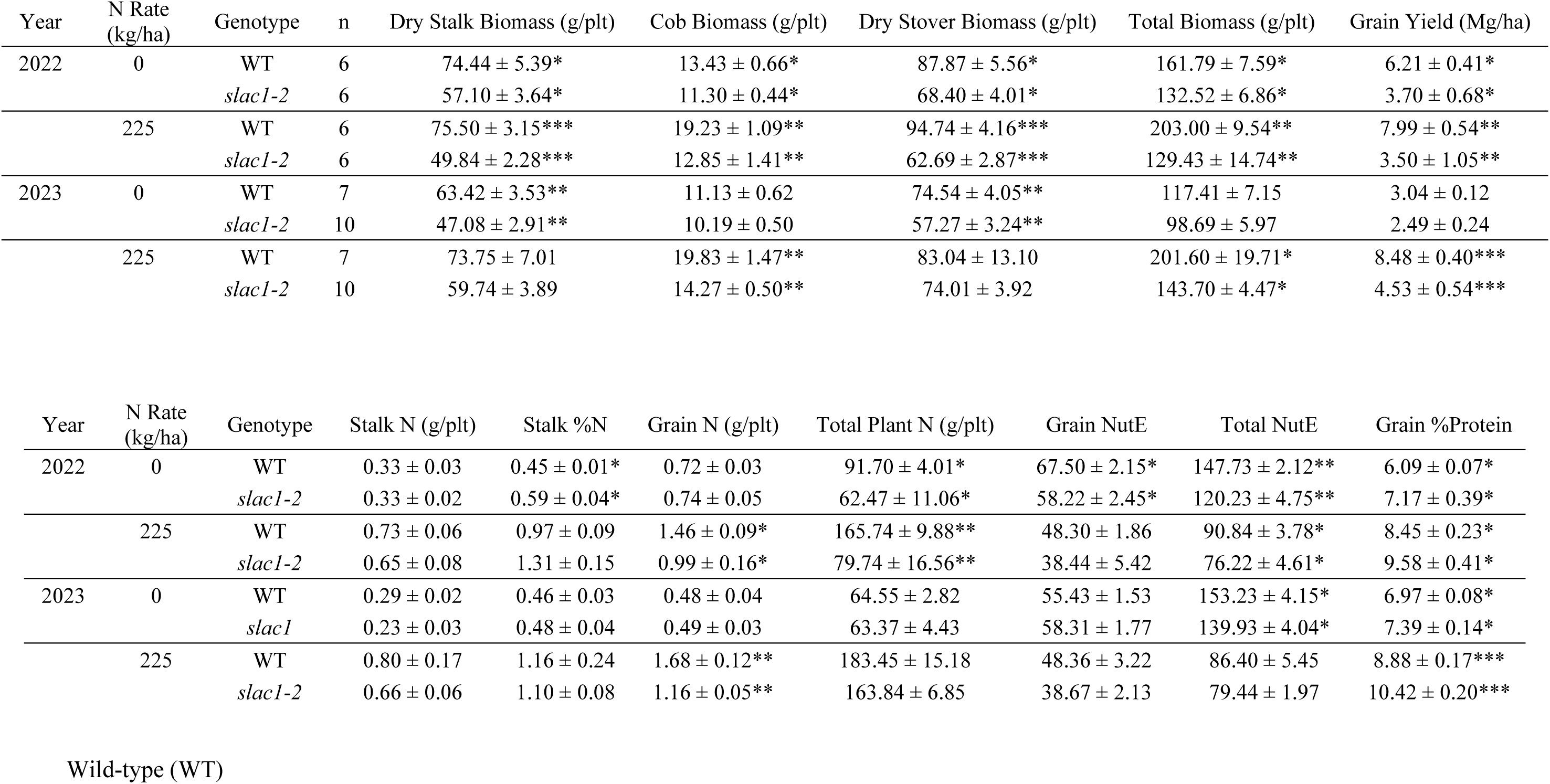
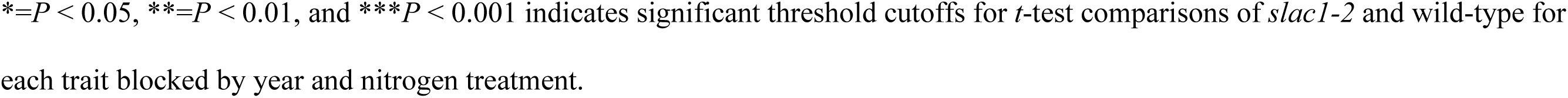
Measured traits from R6 sampling during the 2022 and 2023 Illinois location 2 field trial.

## Discussion

During this study, *slac1-2* inbred and hybrid lines were evaluated in multiple field locations. These trials improved our understanding of the relationships between physiological traits and agronomic performance. The collection of leaf gas exchange, grain yield, and nitrogen utilization data from hybrids with increased *g_s_* and lack of dynamic stomatal response further elucidates the role stomata play in the delicate balance between improved *WUE* and plant productivity.

Thermal imaging has become a well-established high-throughput method for screening diverse populations and predicting crop productivity under varying water conditions (Liu *et al*. 2011; Pignon *et al*., 2021; Wen *et al*., 2023). Significant correlations have been observed between leaf temperature and *g*_s_ using thermal imaging (Winterhalter *et al*., 2011; Zia *et al*., 2013). Thermal imaging was previously used to identify *At*slac1-2, an allelic variant of the original *At*slac1-1, in an EMS-mutagenized *A. thaliana* population (Negi *et al*., 2008). Other studies have observed that *slac1* significantly alters leaf temperature in *A. thaliana*, *O. sativa*, and *Z. mays*; however, all of the thermal imaging data has been collected in controlled growing environments (Kusumi *et al*., 2012; Qi *et al*., 2018; Wang *et al*., 2022). To test these findings in a natural and dynamic environment, field grown *slac1-2 Z. mays* was imaged using a thermal camera. Field grown *slac1-2* was found to have a significantly cooler leaf canopy by an average of 1.25°C compared to wild-type (Fig. 1). This result supports that *slac1-2* has higher *g*_s_ rates in a field environment and at a level that impacts canopy dynamics.

During CO_2_ response measurements, field grown *slac1-2* inbreds were insensitive to a CO_2_ change from 400ppm to 800ppm. These findings are consistent with past results showing *slac1-2* insensitivity to increased CO_2_ measured during a shorter time-course in a controlled growth environment (Qi *et al*., 2018). These findings provide strong evidence that *Z. mays slac1* facilitates anion efflux out of guard cells and controls stomatal closure, similar to *A. thaliana* (Vahisalu *et al*., 2008; Cotelle and Leonhardt, 2016). Unique to this study is the measurement of *Z. mays slac1-2* mutant plant’s response to decreasing CO_2_. An increase in *g*_s_ was observed in *slac1-2*, indicating maintained ability to accumulate anions within the guard cells and open its stomata in response to decreased CO_2_ (Fig. 2A). It is believed that stomatal opening occurs while S-type anions are inactivated and they do not play a vital role (Schwarts *et al*., 1995; Marten *et al*., 2007). However, past evidence would suggest *slac1* can affect the rate of stomatal opening, indicating potential crosstalk between anion and uptake channels (Laanemets *et al*., 2013; Hedrich and Geiger, 2017). The vast difference in *g*_s_ starting values within this data set makes comparing stomatal response rate to decreasing CO_2_ unreliable. Therefore, further studies are needed to determine if *slac1* can alter stomatal opening rates in *Z. mays*.

During three of the *slac1-2* hybrid field trials, survey measurements were collected to determine leaf gas exchange of field adapted plants. These measurements allowed us to capture how *slac1-2* was performing in a dynamic field environment rather than a controlled environment where plants are allowed time to reach steady state. In the 2022 Illinois location 1 field, *slac1-2* showed a significantly higher *g*_s_, *E*, and *C*_i_. However, during the 2023 and 2024 field trials *slac1-2* performed equal to wild-type and no differences were observed (Fig. 3). Similar trends were observed in greenhouse grown rice where *slac1 g*_s_ was only significantly higher than wild-type for a proportion of the developmental timeline (Kusumi *et al*., 2017). Previous data collected from acclimated plants implies that *slac1*-2 shows a consistently more open stomatal phenotype and higher rates of *g*_s_ all the time (Fig. 2A). However, this survey data suggests that in a field environment, *slac1-2* loses its ability to dynamically respond to environmental changes. This can further be supported by the 2024 Arizona survey measurements taken during the weather event that resulted in variable VPD (Fig. 5). Wild-type plants close their stomata and reduce *g*_s_ in response to increased cloud cover and decreased light and VPD. A slight decrease in *g*_s_ can also be observed in *slac1-2* due to a decrease in transpirational demand (lower VPD). However, the absence of a stomatal closure response to increased cloud cover results in a significantly higher *g*_s_ in *slac1-2* compared to wild-type at 9:00. Interestingly, *slac1-2* and wild-type hybrids do not show significantly different *g*_s_ values before and after the light and VPD drop occurred. Thus, because the true phenotype of a *slac1* mutant is lack of response, depending on the environment, mutants could have either a higher or lower *g*_s_ value than wild-type plants when instantaneous measurements are taken.

Leaf stable carbon isotope composition (δ^13^C_leaf_) was measured for tissue collected from greenhouse and field grown *slac1-2* plants. Assaying δ^13^C_leaf_ has been shown to be an integrated measure of carbon metabolism linked to *iWUE* in C_4_ species (O’Leary, 1981; Twohey III *et al*., 2019). Based on C_4_ isotope theory, increased *g*_s_ would result in a less negative δ^13^C (O’Leary, 1988), which has been observed in other *Z. mays* studies (Avramova *et al*., 2019; Crawford *et al*., 2024). In the greenhouse and field grown *slac1-2* plants, significantly less negative δ^13^C values were observed compared to wild-type (*P* < 0.01 and *P* < 0.001 respectively, *t*-test, Table 1). These results indicate that *slac1-2* stomatal hyposensitivity is altering plant *g*_s_ not only at the time of extreme environmental changes, but throughout plant development. Furthermore, because of the less negative δ^13^C value, even though not all instantaneous gas exchange measurements capture greater *g_s_* in *slac1* mutant plants, the cumulative effect of the mutation is higher *g_s_* and greater water loss.

To determine if stomatal limitation of CO_2_ uptake for photosynthesis exists in C_4_ Z. *mays*, *A*/*C*_i_ curves were measured. During the *A*/*C*_i_ curves no significant difference was observed in *A*_max_, although *slac1-2* mutants tended to be slightly lower. During increased *C*_a_ (800ppm), *slac1-2* had a significantly higher *C*_i_/*C*_a_ value compared to wild-type, consistent with a greater stomatal aperture and inability to close in response to elevated CO_2_ concentrations (Fig. 4). Observing higher *C*_i_/*C*_a_ values with no significant benefit in *A*_net_ supports the idea that C_4_ often remains CO_2_ saturated and increased *g*_s_ does not increase productivity. Although not shown to be significant, the opposite trend was observed in rice, a C_3_ species. Two studies observed slight increases in *A*_net_ under high *C*_i_/*C*_a_ levels in *slac1-2* compared to wild-type (Kusumi *et al*., 2012; Kusumi *et al*., 2017).

To further determine if *Z. mays* productivity is at all limited by CO_2_ uptake, we collected grain yield data from seven different field environments. Some significant differences were seen between *slac1-2* and wild-type hybrids; however, *slac1-2* never out yielded wild-type (Fig. 6). These results highlight the utility of the C_4_ carbon concentrating mechanism, and the extent to which it enables a reduced *g*_s_ without a corresponding decrease in *A*_net_ or grain yield. The diverse environments highlight the performance of a CO_2_ concentrating mechanism in wild-type hybrids under well-watered and high light environments. Therefore, an opportunity exists for targeting *g*_s_ in *Z. mays* to improve *iWUE* while maintaining plant productivity. These results are consistent with a previous study targeting *slac1* in a field environment to determine its effects on yield under drought conditions. *Z. mays* inbred and hybrid lines overexpressing the protein kinases ZmCPK35 and ZmCPK37 increased yields by activating *slac1* and inducing stomatal closure under drought conditions (Li *et al*., 2022).

There is mounting evidence that plant transpiration passively regulates nitrogen uptake (Mengel and Kirby, 1987; Novák and Vidovič, 2003; Niu *et al*., 2007; Kunrath *et al*., 2020). The *slac1-2* field trials allowed us to directly measure if increased *g*_s_ would lead to higher plant nitrogen uptake or utilization. Despite *slac1-2* inbreds having a higher leaf %N at V10-12 (Table 1), a greater %N was not observed in the hybrid R6 data (Table 2). During the two field trials both *slac1-2* and wild-type had equally reduced grain nitrogen under a 0 kg/ha nitrogen rate, but wild-type had a significantly greater response to a nitrogen rate of 225 kg/ha compared to *slac1-2*. Overall, nitrogen availability seemed to be similar in *slac1-2* and wild-type; however, *slac1-2* did not utilize the available nitrogen. This can be observed in the grain nitrogen utilization efficiency (NutE) values where *slac1-2* was significantly lower than wild-type in almost all conditions (Table 2).

The *slac1-2* hybrid facilitated the investigation of stomatal dynamics in a *Z. mays* background comparable to commercially relevant germplasm and in production field environments. Using gas exchange, grain yield, and nitrogen data to measure *slac1-2,* we observed that the CO_2_ concentrating mechanism in *Z. mays* is not significantly limited by *g*_s_ under a variety of field conditions. This is supported by the lack of *A*_net_ or grain yield increases in *slac1-2* hybrids across all field trials. Future studies using an overexpression of *slac1-2* might allow us to reduce *g*_s_ and identify an optimum response sensitivity that maximizes *WUE* without negatively affecting *A*_net_ and grain yield. Additionally, because *slac1-2* removes stomata as the major regulator of plant water loss, alternate mechanisms that modulate plant water potential can be probed to reveal novel approaches to further improving *WUE*.

## Acknowledgments and Funding

We thank all technical support and field managers from the Studer, Moose, Gore, and Pauli Labs. Much thanks to Andrew Leaky for the use of the stomatal imaging microscope. We would like to thank the Maize Genetic Cooperation Stock Center for supplying the original *slac1-2* germplasm, especially Marty Sachs for valuable discussion. This research was performed with support from the Center for Research on Programmable Plant Systems under National Science Foundation Grant No. DBI-2019674 and the United States Department of Agriculture Hatch funds as well as additional funding to D. P. (NSF IOS 2102120; NSF IOS 2023310; USDA NIFA SCRI # 2021-51181-35903; Cotton Inc. 23-890).

## Author Contributions

R.J.T and A.J.S conceived the project. R.J.T. and A.J.S. developed the germplasm. R.J.T, A.J.S, S.P.M, M.A.G, and D.P designed field experiments. R.J.T performed inbred field experiments. R.J.T, C.G.C, C.L, H.B, S.C, B.P, M.H, and L.W performed hybrid field experiments. R.J.T analyzed the data. R.J.T and A.J.S wrote the manuscript. All authors reviewed and revised the manuscript.

We have no conflicts of interest to disclose.

## Notes

### Competing Interest Statement

The authors have declared no competing interest.

## Citations

Ainsworth EA, Long SP (2021) 30 years of free-air carbon dioxide enrichment (FACE): What have we learned about future crop productivity and its potential for adaptation? Global change biology 27: 27–49

Assmann SM, Jegla T (2016) Guard cell sensory systems: recent insights on stomatal responses to light, abscisic acid, and CO2. Current opinion in plant biology 33: 157–167

Avramova V, Meziane A, Bauer E, Blankenagel S, Eggels S, Gresset S, Grill E, Niculaes C, Ouzunova M, Poppenberger B (2019) Carbon isotope composition, water use efficiency, and drought sensitivity are controlled by a common genomic segment in maize. Theoretical and Applied Genetics 132: 53–63

Cheng C-Y, Li Y, Varala K, Bubert J, Huang J, Kim GJ, Halim J, Arp J, Shih H-JS, Levinson G (2021) Evolutionarily informed machine learning enhances the power of predictive gene-to-phenotype relationships. Nature communications 12: 5627

Christian JI, Basara JB, Hunt ED, Otkin JA, Furtado JC, Mishra V, Xiao X, Randall RM (2021) Global distribution, trends, and drivers of flash drought occurrence. Nature communications 12: 6330

Copernicus Climate Change Service, 2024, https://climate.copernicus.eu/global-climate-highlights-2024

Cotelle V, Leonhardt N (2016) 14-3-3 proteins in guard cell signaling. Frontiers in plant science 6: 1210

Crawford JD, Twohey Iii RJ, Pathare VS, Studer AJ, Cousins AB (2024) Differences in stomatal sensitivity to CO2 and light influence variation in water use efficiency and leaf carbon isotope composition in two genotypes of the C4 plant Zea mays. Journal of experimental botany 75: 6748–6761

European State of the Climate 2023 (ESOTC)" published by the Copernicus Climate Change Service (C3S)

Farquhar GD, Sharkey TD (1982) Stomatal conductance and photosynthesis. Annual review of plant physiology 33: 317–345

Gamelin BL, Feinstein J, Wang J, Bessac J, Yan E, Kotamarthi VR (2022) Projected US drought extremes through the twenty-first century with vapor pressure deficit. Scientific Reports 12: 8615

Geiger D, Scherzer S, Mumm P, Stange A, Marten I, Bauer H, Ache P, Matschi S, Liese A, Al-Rasheid KAS (2009) Activity of guard cell anion channel SLAC1 is controlled by drought-stress signaling kinase-phosphatase pair. Proceedings of the National Academy of Sciences 106: 21425–21430

Goodstein DM, Shu S, Howson R, Neupane R, Hayes RD, Fazo J, Mitros T, Dirks W, Hellsten U, Putnam N (2012) Phytozome: a comparative platform for green plant genomics. Nucleic acids research 40: D1178–D1186

Guo Y, Shi Y, Wang Y, Liu F, Li Z, Qi J, Wang Y, Zhang J, Yang S, Wang Y (2023) The clade F PP2C phosphatase ZmPP84 negatively regulates drought tolerance by repressing stomatal closure in maize. New Phytologist 237: 1728–1744

He X, Wang Y, Munawar A, Zhu J, Zhong J, Zhang Y, Guo H, Zhu Z, Baldwin IT, Zhou W (2025) Manipulating stomatal aperture by silencing StSLAC1 affects potato plant–herbivore– parasitoid tritrophic interactions under drought stress. New Phytologist

Hedrich R, Geiger D (2017) Biology of SLAC 1-type anion channels–from nutrient uptake to stomatal closure. New Phytologist 216: 46–61

Keller BU, Hedrich R, Raschke K (1989) Voltage-dependent anion channels in the plasma membrane of guard cells. Nature 341: 450–453

Kolbe AR, Brutnell TP, Cousins AB, Studer AJ (2018) Carbonic anhydrase mutants in Zea mays have altered stomatal responses to environmental signals. Plant Physiology 177: 980–989

Kunrath TR, Lemaire G, Teixeira E, Brown HE, Ciampitti IA, Sadras VO (2020) Allometric relationships between nitrogen uptake and transpiration to untangle interactions between nitrogen supply and drought in maize and sorghum. European Journal of Agronomy 120: 126145

Kusumi K, Hashimura A, Yamamoto Y, Negi J, Iba K (2017) Contribution of the S-type anion channel SLAC1 to stomatal control and its dependence on developmental stage in rice. Plant and Cell Physiology 58: 2085–2094

Kusumi K, Hirotsuka S, Kumamaru T, Iba K (2012) Increased leaf photosynthesis caused by elevated stomatal conductance in a rice mutant deficient in SLAC1, a guard cell anion channel protein. Journal of experimental botany 63: 5635–5644

Laanemets K, Wang YF, Lindgren O, Wu J, Nishimura N, Lee S, Caddell D, Merilo E, Brosche M, Kilk K (2013) Mutations in the SLAC 1 anion channel slow stomatal opening and severely reduce K+ uptake channel activity via enhanced cytosolic [Ca2+] and increased Ca2+ sensitivity of K+ uptake channels. New Phytologist 197: 88–98

Lawson T, Blatt MR (2014) Stomatal size, speed, and responsiveness impact on photosynthesis and water use efficiency. Plant physiology 164: 1556–1570

Lawson T, Vialet-Chabrand S (2019) Speedy stomata, photosynthesis and plant water use efficiency. New Phytologist 221: 93–98

Lawson T, von Caemmerer S, Baroli I (2011) Photosynthesis and stomatal behaviour. Progress in botany 72: 265–304

Leakey ADB, Ferguson JN, Pignon CP, Wu A, Jin Z, Hammer GL, Lobell DB (2019) Water use efficiency as a constraint and target for improving the resilience and productivity of C3 and C4 crops. Annual review of plant biology 70: 781–808

Lee H, Calvin K, Dasgupta D, Krinner G, Mukherji A, Thorne P, Trisos C, Romero J, Aldunce P, Barret K (2023) IPCC, 2023: Climate Change 2023: Synthesis Report, Summary for Policymakers. Contribution of Working Groups I, II and III to the Sixth Assessment Report of the Intergovernmental Panel on Climate Change [Core Writing Team, H. Lee and J. Romero (eds.)]. IPCC, Geneva, Switzerland.

Lee H, Calvin K, Dasgupta D, Krinner G, Mukherji A, Thorne P, Trisos C, Romero J, Aldunce P, Barrett K (2023) Climate change 2023: synthesis report. Contribution of working groups I, II and III to the sixth assessment report of the intergovernmental panel on climate change. The Australian National University

Lee SC, Lan W, Buchanan BB, Luan S (2009) A protein kinase-phosphatase pair interacts with an ion channel to regulate ABA signaling in plant guard cells. Proceedings of the National Academy of Sciences 106: 21419–21424

Li XD, Gao YQ, Wu WH, Chen LM, Wang Y (2022) Two calcium-dependent protein kinases enhance maize drought tolerance by activating anion channel ZmSLAC1 in guard cells. Plant Biotechnology Journal 20: 143–157

Liu X, Mak M, Babla M, Wang F, Chen G, Veljanoski F, Wang G, Shabala S, Zhou M, Chen Z- H (2014) Linking stomatal traits and expression of slow anion channel genes HvSLAH1 and HvSLAC1 with grain yield for increasing salinity tolerance in barley. Frontiers in plant science 5: 634

Liu Y, Subhash C, Yan J, Song C, Zhao J, Li J (2011) Maize leaf temperature responses to drought: Thermal imaging and quantitative trait loci (QTL) mapping. Environmental and Experimental Botany 71: 158–165

López J, Way DA, Sadok W (2021) Systemic effects of rising atmospheric vapor pressure deficit on plant physiology and productivity. Global Change Biology 27: 1704–1720

Mansfield TA (1983) Movements of stomata. Science Progress (1933-): 519–542

Marten H, Konrad KR, Dietrich P, Roelfsema MRG, Hedrich R (2007) Ca2+-dependent and- independent abscisic acid activation of plasma membrane anion channels in guard cells of Nicotiana tabacum. Plant Physiology 143: 28–37

McDaniel RL, Munster C, Nielsen-Gammon J (2017) Crop and location specific agricultural drought quantification: part III. Forecasting water stress and yield trends. Transactions of the ASABE 60: 741–752

McDonald RI, Girvetz EH (2013) Two challenges for US irrigation due to climate change: increasing irrigated area in wet states and increasing irrigation rates in dry states. PloS one 8: e65589

Mengel K, Kirkby EA (1987) Principles of plant nutrition,

Negi J, Matsuda O, Nagasawa T, Oba Y, Takahashi H, Kawai-Yamada M, Uchimiya H, Hashimoto M, Iba K (2008) CO2 regulator SLAC1 and its homologues are essential for anion homeostasis in plant cells. Nature 452: 483–486

Niu J, Chen F, Mi G, Li C, Zhang F (2007) Transpiration, and nitrogen uptake and flow in two maize (Zea mays L.) inbred lines as affected by nitrogen supply. Annals of Botany 99: 153–160

Novák V, Vidovič J (2003) Transpiration and nutrient uptake dynamics in maize (Zea mays L.). Ecological Modelling 166: 99–107

Novick KA, Ficklin DL, Grossiord C, Konings AG, Martínez-Vilalta J, Sadok W, Trugman AT, Williams AP, Wright AJ, Abatzoglou JT (2024) The impacts of rising vapour pressure deficit in natural and managed ecosystems. Plant, Cell & Environment

O’Leary MH (1981) Carbon isotope fractionation in plants. Phytochemistry 20: 553-567 O’Leary MH (1988) Carbon isotopes in photosynthesis. Bioscience 38: 328–336

Pei Z-M, Kuchitsu K, Ward JM, Schwarz M, Schroeder JI (1997) Differential abscisic acid regulation of guard cell slow anion channels in Arabidopsis wild-type and abi1 and abi2 mutants. The Plant Cell 9: 409–423

Pei Z-M, Murata Y, Benning G, Thomine S, Klüsener B, Allen GJ, Grill E, Schroeder JI (2000) Calcium channels activated by hydrogen peroxide mediate abscisic acid signalling in guard cells. Nature 406: 731–734

Pignon CP, Fernandes SB, Valluru R, Bandillo N, Lozano R, Buckler E, Gore MA, Long SP, Brown PJ, Leakey ADB (2021) Phenotyping stomatal closure by thermal imaging for GWAS and TWAS of water use efficiency-related genes. Plant physiology 187: 2544–2562

Portwood JL, Woodhouse MR, Cannon EK, Gardiner JM, Harper LC, Schaeffer ML, Walsh JR, Sen TZ, Cho KT, Schott DA (2019) MaizeGDB 2018: the maize multi-genome genetics and genomics database. Nucleic acids research 47: D1146–D1154

Qi G-N, Yao F-Y, Ren H-M, Sun S-J, Tan Y-Q, Zhang Z-C, Qiu B-S, Wang Y-F (2018) The S-type anion channel ZmSLAC1 plays essential roles in stomatal closure by mediating nitrate efflux in maize. Plant and Cell Physiology 59: 614–623

R Core Team (2021). R: A language and environment for statistical computing. R Foundation for Statistical Computing, Vienna, Austria. URL https://www.R-project.org/.

Ren H, Su Q, Hussain J, Tang S, Song W, Sun Y, Liu H, Qi G (2021) Slow anion channel GhSLAC1 is essential for stomatal closure in response to drought stress in cotton. Journal of Plant Physiology 258: 153360

Roelfsema MRG, Hedrich R (2005) In the light of stomatal opening: new insights into ‘the Watergate’. New Phytologist 167: 665–691

Schwartz A, Ilan N, Schwarz M, Scheaffer J, Assmann SM, Schroeder JI (1995) Anion-channel blockers inhibit S-type anion channels and abscisic acid responses in guard cells. Plant physiology 109: 651–658

Settles AM, Latshaw S, McCarty DR (2004) Molecular analysis of high-copy insertion sites in maize. Nucleic acids research 32: e54–e54

Twohey III RJ, Roberts LM, Studer AJ (2019) Leaf stable carbon isotope composition reflects transpiration efficiency in Zea mays. The Plant Journal 97: 475–484

Vahisalu T, Kollist H, Wang Y-F, Nishimura N, Chan W-Y, Valerio G, Lamminmäki A, Brosché M, Moldau H, Desikan R (2008) SLAC1 is required for plant guard cell S-type anion channel function in stomatal signalling. Nature 452: 487–491

von Caemmerer S, Evans JR (2010) Enhancing C3 photosynthesis. Plant Physiology 154: 589–592

Wallace JS (2000) Increasing agricultural water use efficiency to meet future food production. Agriculture, ecosystems & environment 82: 105–119

Wang Z, Ouyang Y, Ren H, Wang S, Xu D, Xin Y, Hussain J, Qi G (2022) Transcriptome profiling of Arabidopsis slac1-3 mutant reveals compensatory alterations in gene expression underlying defective stomatal closure. Frontiers in Plant Science 13: 987606

Wen T, Li J-H, Wang Q, Gao Y-Y, Hao G-F, Song B-A (2023) Thermal imaging: The digital eye facilitates high-throughput phenotyping traits of plant growth and stress responses. Science of The Total Environment: 165626

Winterhalter L, Mistele B, Jampatong S, Schmidhalter U (2011) High throughput phenotyping of canopy water mass and canopy temperature in well-watered and drought stressed tropical maize hybrids in the vegetative stage. European Journal of Agronomy 35: 22–32

Wong SC, Cowan IR, Farquhar GD (1979) Stomatal conductance correlates with photosynthetic capacity. Nature 282: 424–426

Zia S, Romano G, Spreer W, Sanchez C, Cairns J, Araus JL, Müller J (2013) Infrared thermal imaging as a rapid tool for identifying water-stress tolerant maize genotypes of different phenology. Journal of Agronomy and Crop Science 199: 75–84

