## Supplemental Data for "Investigating the role of stomatal dynamics on agronomic traits using a *slac1-2 Zea mays* mutant"

**Supplemental Figures**

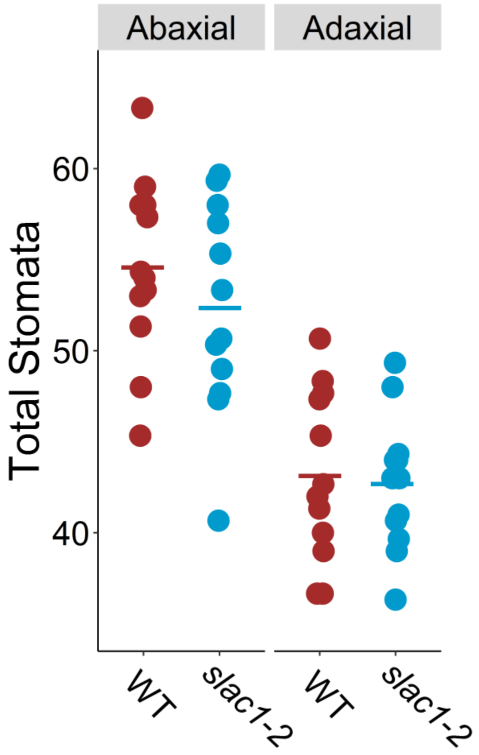


**Sup. Figure 1**

Total stomata per 0.8mm x 0.8mm image of the abaxial and adaxial leaf surfaces are shown for *slac1-2* (blue) and its comparative wild-type (WT, red) in a W22 inbred background. Points represent individual replicates, and the solid dash represents average total stomata. No significant differences were seen between the two genotypes ($P > 0.05$, *t*-test).

### Supplemental Tables

Sup. Table 1

| | Stomatal conductance ($g_s$) mol $m^{-2}$ $s^{-1}$ | | | Net Photosynthesis (A) umol $m^{-2}$ $s^{-1}$ | | | Transpiration (E) mol $m^{-2}$ $s^{-1}$ | | |
|---|---|---|---|---|---|---|---|---|---|
| | 400 ppm* | 800 ppm* | 100 ppm | 400 ppm | 800 ppm | 100 ppm | 400 ppm* | 800 ppm* | 100 ppm |
| *slac1-2* | 0.418±0.017 | 0.398±0.0189 | 0.476±0.0180 | 40.320±1.283 | 42.629±0.893 | 15.394±0.750 | 0.00619±0.000224 | 0.00594±0.000269 | 0.00688±0.000258 |
| WT | 0.330±0.0127 | 0.156±0.00438 | 0.485±0.0166 | 40.660±0.786 | 42.924±0.797 | 15.656±0.562 | 0.00501±0.000176 | 0.00248±0.0000625 | 0.00714±0.000203 |

Wild-type (WT)

* Significant difference between *slac1-2* and WT ($P < 0.01$, *t*-test)

Sup. Table 2

**R6 2023 Phenotypes**

| N Rate (kg/ha) | Genotype | n | DTA | DTS | Plant Height (cm) |
|---|---|---|---|---|---|
| 0 | WT Hyb | 7 | 1313.00 ± NA | 1402.00 ± NA | 240.48 ± 1.51** |
| | *slac1-2* Hyb | 10 | 1402.00 ± 0.00 | 1480.00 ± 7.60 | 212.89 ± 3.38** |
| 225 | WT Hyb | 7 | 1268.00 ± 0.00 | 1325.00 ± 6.41 | 250.61 ± 1.62* |
| | *slac1-2* Hyb | 10 | 1369.75 ± 4.81 | 1391.25 ± 4.81 | 238.78 ± 1.89* |

Wild-type (WT)

*= $P < 0.01$

**= $P < 0.001$
